# Glutamatergic excitation suppresses latent cortical pacemaking

**DOI:** 10.64898/2026.10.26.759671

**Authors:** Daniil P. Aksenov, Evan Doubovikov, Natalya A. Serdyukova, Mohammed M. Minhaj, Steven B. Greenberg

**Affiliations:** Endeavor Health, Evanston, USA; Pritzker School of Medicine, Chicago, USA; Department of Biomedical Engineering, Northwestern University, Evanston, USA

## Abstract

Glutamate provides the principal excitatory drive in the cortex, so reducing ionotropic glutamatergic transmission is expected to suppress neuronal activity. We asked whether reducing ongoing glutamatergic excitation can reveal a previously unrecognized form of neuronal activity: a local population timing mechanism. Combining two established single-neuron constraints (near-threshold susceptibility and spike-history-dependent loss of excitability) we proposed an availability reservoir: a local population can generate a burst only when enough neurons are both susceptible and recovered. This hypothesis predicted a counterintuitive result: local glutamate-receptor antagonism should not simply silence the awake cortex, but should allow the eligible pool to refill and reveal recurrent burst-quiescence. In the awake rabbit cortex, local glutamate-receptor antagonism produced stereotyped local-field-potential and multi-unit population bursts separated by quiescence. Increasing GABAergic inhibition progressively reduced burst size and recurrence before abolishing the pacemaker regime, whereas reducing GABAergic inhibition under baseline conditions produced epileptiform activity with different timing and intra-burst structure. Thus, glutamatergic excitation can mask a recovery-dependent cortical population pacemaker regime. The same framework distinguishes recovery-dependent burst-quiescence from disinhibited epileptiform bursting, providing an operational separation between visually similar but mechanistically different brain states.

## Introduction

In the neocortex, excitation is treated mainly as the driver of activity. We asked whether ongoing excitation can also mask the timing mechanism that determines when a coordinated population burst can occur. After neurons fire, many become temporarily harder to recruit again. If glutamatergic input keeps recruiting scattered neurons before a sufficiently large group has recovered, the cortex may remain active but fail to generate an organized population burst. This creates the central possibility tested here: reducing excitation may reveal burst timing by allowing a recovered population to accumulate.

The burst–quiescence state revealed by reduced glutamatergic excitation represents a previously unrecognized form of cortical population activity organized by local population timing. This raised the possibility that the state reflects latent population pacemaking. Pacemaking has a long-standing neurophysiological meaning beyond cardiac or respiratory control. Early experiments showed that isolated somata of *Aplysia* pacemaker neurons continued to discharge spontaneously without synaptic input.^1^ The term also describes neuronal sources of hippocampal oscillations,^2^ while the excitable-media framework extends it to self-organized firing clusters rather than requiring intrinsically oscillating cells.^3^ Thus, we use “pacemaker” operationally and at the population level. A cortical population pacemaker regime is a state of excitable cortex in which recurrent population bursts emerge after recovery-dependent quiescence because a susceptible and recovered local population crosses a threshold for recruitment. This usage does not imply a dedicated cortical pacemaker cell, a fixed anatomical pacemaker nucleus or a deterministic clock. It is consistent with the excitable-media framework, in which pacemakers can emerge as self-organized firing clusters rather than only as groups of limit-cycle oscillators.^3^

The hypothesis began by combining two established single-neuron observations that had not previously been connected as a population pacemaker mechanism. First, neocortical pyramidal neurons can enter near-threshold susceptible states and generate rhythmic firing when held close to spike threshold.^4^ Second, recent spiking can elevate pyramidal-cell spike threshold in vivo, imposing a slow recovery constraint lasting hundreds of milliseconds to about a second.^5^ Neither observation proposed a cortical availability reservoir. The reservoir hypothesis is the inference introduced here from their combination. We formalized this idea as an availability–recruitment model in which recovery refills an eligible population after each burst (Supplementary Methods 4).

The inference is restrictive. A neuron can participate in a population burst only when it is both susceptible and sufficiently recovered. We call the fraction of neurons satisfying both conditions the availability reservoir. A burst discharges this reservoir by recruiting many neurons into recovery; quiescence refills it. A new burst becomes possible when recovered susceptible tissue is large enough for local recruitment to cross threshold. Timing is therefore not imposed by a clock-like cortical oscillator, but by recovery-dependent refill followed by thresholded recruitment.

This framing reverses the usual role assigned to glutamatergic excitation. Under normal awake conditions, ongoing glutamatergic recruitment repeatedly activates neurons and repeatedly resets recovery. The cortex can therefore contain susceptible states in principle while failing to express them as population timing. We predicted that reducing glutamatergic recruitment would not necessarily silence cortex; it could allow recovery to proceed broadly, refill the availability reservoir and reveal a latent pacemaker regime. Here, latent means that the cellular conditions required for the regime exist before perturbation, whereas ongoing glutamatergic recruitment prevents their expression as population pacemaking.

Here we report a focused sequence of tests of this prediction. First, we formulate the reservoir hypothesis without relying on an anatomical pacemaker site. Second, we test the prediction that local glutamate-receptor antagonism reveals burst-quiescence in awake cortex. Third, we characterize the physiology of the pacemaking bursts. Fourth, in complementary experiments, we tested whether increasing GABA-A-receptor signalling suppresses pacemaking and whether reducing it produces a distinct disinhibited regime. This sequence separates recovery-dependent pacemaking from disinhibited epileptiform activity.

## Results

### A reservoir hypothesis predicted pacemaking

The conceptual model separated two components formalized in Supplementary Methods 4: an availability variable describing recovery after recent activity, and a probabilistic initiation process describing when recovered tissue generated the next population event. Ongoing glutamatergic activity repeatedly depleted availability, whereas reducing glutamatergic recruitment allowed availability to rebuild.

The conceptual model predicted burst-quiescence after glutamate-receptor antagonism. Recovery produced a short low-probability interval after each event, after which burst initiation followed the phenomenological stochastic initiation process defined in Supplementary Methods 4. We next asked whether the same qualitative transition emerged in a physiological two-compartment cortical model. Removing glutamate-mediated recruitment converted high-frequency resting activity into low-frequency compound bursting at approximately 0.5 Hz in simulated local-field-potential and multi-unit signals (Fig. 1a,b). Thus, both models predicted that glutamate-receptor antagonism would reveal recurrent bursting rather than simply silence cortex.

**Figure 1.**
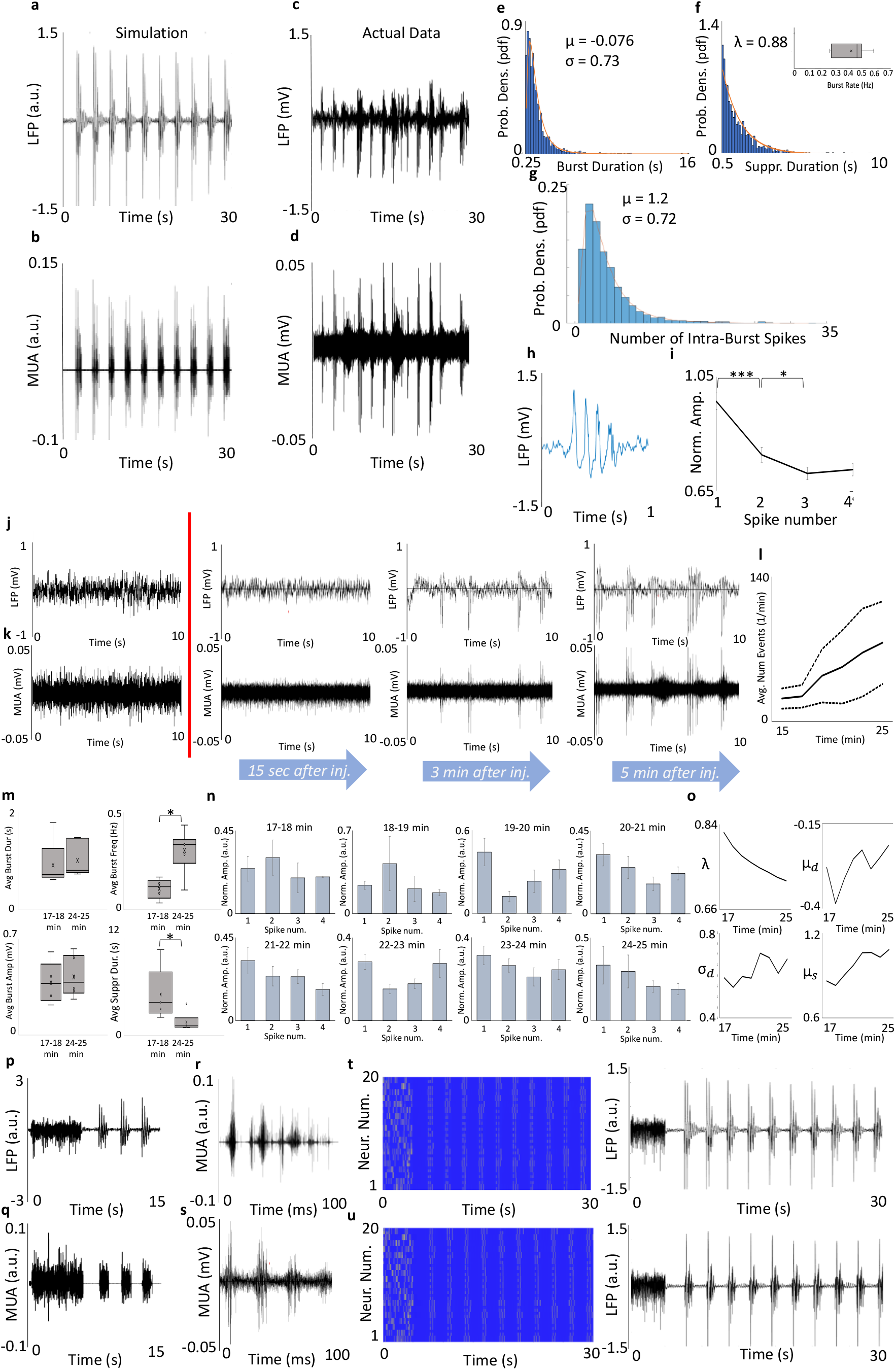
Glutamate antagonism reveals the predicted cortical pacemaker regime. a,b, Physiological simulations predicted recurring LFP and MUA bursts after removal of glutamate-mediated recruitment; glutamate removal produced low-frequency compound bursts at approximately 0.5 Hz. c,d, Representative awake-cortex recordings after local DGG show the corresponding burst-quiescence pattern. e-g, Distributions characterizing the pacemaker regime in DGG experiments: burst duration fitted by a log-normal distribution, suppression duration fitted by an exponential distribution and intra-burst LFP peak number fitted by a log-normal distribution. The stable pacemaker burst rate was approximately 0.4 Hz. h,i, Example pacemaking burst and average cascading profile. j,k, LFP and MUA examples during the emergence phase after injection. l-o, Time-dependent evolution of event number, burst parameters, intra-burst profile and fitted distribution parameters. p-s, Long-duration model/data comparisons and MUA substructure within compound LFP events; model and experimental MUA show that a single compound LFP peak can contain multiple sub-population recruitment events over approximately 100 ms. t,u, Simulations with and without extracellular-potential feedback show its contribution to burst synchrony and amplitude. The asterisks *,**,***, denote a significance level of at least 5%, 1%, and 0.1% respectively.

The physiological model also tested the contribution of extracellular-potential feedback to population recruitment. Repeating the simulation without calculating the extracellular field reduced burst synchrony past the initial peak (Fig. 1t,u), indicating that extracellular fields can facilitate recruitment under simulated glutamate antagonism. These simulations did not independently identify the process that determines the inter-burst interval.

### Glutamate antagonism reveals the predicted pacemaker regime

We tested the prediction in awake rabbit somatosensory cortex using local gamma-D-glutamylglycine (DGG), a broad ionotropic glutamate-receptor antagonist. In the DGG-only cohort, recordings were obtained from six rabbits. DGG was applied specifically to test whether glutamate-receptor antagonism would reveal, rather than suppress, recurrent burst-quiescence. Before injection, the same sites did not show a comparable recurrent burst-quiescence state.

After DGG, local-field-potential and multi-unit recordings developed stereotyped population bursts separated by prolonged intervals of reduced activity (Fig. 1c,d,j,k,p,q). Vehicle injections did not produce the burst-quiescence pattern, consistent with our previous reports that equal-volume vehicle injections do not alter LFP or MUA activity.^6–8^ Glutamate-receptor antagonism therefore revealed the predicted pacemaker regime rather than simply suppressing cortical activity.

By the operational definition above, we refer to this state as a population pacemaker regime because it consisted of recurrent, recovery-separated recruitment of local population activity rather than autonomous firing by a specialized cortical cell. The stable pacemaker regime occurred at an average burst rate of approximately 0.4 Hz, close to the approximately 0.5 Hz operating regime predicted by the physiological model (Fig. 1f inset).

The onset of the pacemaking state occurred over minutes, consistent with a pharmacological operating-point transition rather than an externally imposed rhythm. In representative emergence traces, 15 s after DGG the LFP showed higher-frequency activity before stereotyped bursts had stabilized; by 3 min, early bursts appeared with wider and less regular inter-burst intervals; by 5 min, LFP and MUA showed the characteristic burst-quiescence pattern with reduced MUA baseline during suppression periods (Fig. 1j,k). Across experiments, event number, burst duration, burst amplitude, burst frequency (p < 0.05) and suppression duration (p < 0.05) changed across the post-injection period (Fig. 1l,m,o).

### Pacemaking bursts show reservoir physiology

The pacemaker regime revealed a previously undescribed burst physiology in awake cortex. Across the DGG cohort, burst durations were right-skewed and fitted by a log-normal distribution with mu = -0.076 and sigma = 0.73 (Fig. 1e). Suppression durations were fitted by an exponential distribution with lambda = 0.88 (Fig. 1f). The number of multi-unit peaks within bursts was also right-skewed and fitted by a log-normal distribution with mu = 1.2 and sigma = 0.72 (Fig. 1g). Thus, pacemaking bursts were stereotyped population events with variable size and interval statistics. The suppression-duration fit was consistent with a stochastic post-recovery initiation component, without implying that the deterministic availability equation alone generates an exact exponential process.

Multi-unit activity resolved internal burst structure that was only partly visible in the local field potential. A single local-field-potential burst often contained several rapid multi-unit peaks over a short recruitment window (Fig. 1h,r,s). In model and experiment, this MUA substructure occurred on an approximately 100 ms timescale, indicating that a single LFP peak can contain several near-synchronous sub-population recruitment events.

In pacemaking bursts, the mean normalized LFP amplitude declined across successive peaks, producing a cascading profile (Fig. 1i,n). This declining profile is consistent with rapid recruitment followed by depletion of a local eligible pool: the first recruitment wave engages the largest available fraction, and later peaks occur from a progressively smaller remaining pool. The cascading profile was not a generic property of all bursts; it emerged after stabilization of the pacemaking regime (Fig. 1n).

### Muscimol progressively reduces pacemaker burst amplitude and recurrence

We used muscimol to titrate the availability reservoir directly. In the model, a gradual increase in tonic inhibition progressively reduced burst expression, whereas a larger increase terminated activity more rapidly (Fig. 2a-f).

**Figure 2.**
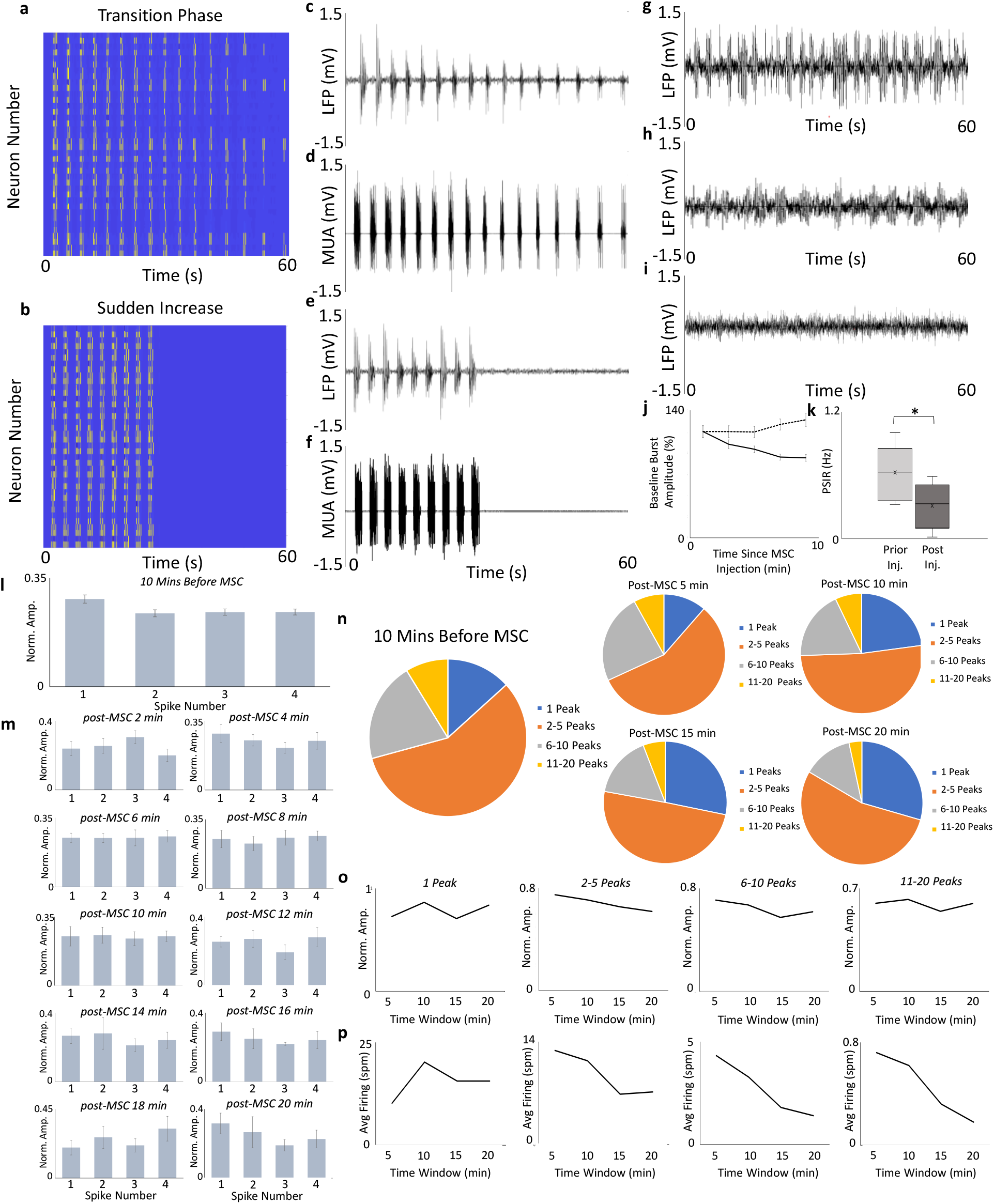
Muscimol progressively reduces pacemaker burst amplitude and recurrence. All experimental times are expressed relative to muscimol injection. a,b, Model raster plots during the transition and quiescent phases produced by increasing tonic GABA conductance. c-f, Simulated LFP and MUA showing progressive weakening and eventual termination of burst activity as inhibition increases. g-i, One-minute experimental LFP segments showing established pacemaking before muscimol, smaller and less frequent bursts 5 min after muscimol, and near-quiescence 10 min after muscimol. j,k, LFP burst amplitude in the DGG + MSC group decreases in comparison to the DGG group (error bars denote standard error) and burst recurrence decrease relative to the established pre-muscimol pacemaker period. l,m, The characteristic cascading intra-burst LFP profile present before muscimol is progressively lost after injection. n, High-peak-count bursts become less frequent relative to simpler events after muscimol. o,p, Intra-burst peak amplitudes and multi-peak event rates decline and reach a plateau across the post-muscimol period.

Before muscimol, recordings showed the established recurrent burst-quiescence pattern. Five minutes after muscimol, LFP bursts were smaller and less frequent; 10 min after muscimol, bursts were nearly absent and the recording approached quiescence (Fig. 2g-i). The amplitude of LFP bursts was significantly lower (p < 0.05) in the DGG+MSC group than in the DGG-only group, after 10 minutes post-injection (Fig 2j). Within the MSC group, the post-suppression initiation rate (p < 0.05) decreased relative to the pre-muscimol pacemaker period (Fig. 2k). The reduction in the post-suppression initiation rate corresponded to progressively longer intervals between successful bursts.

LFP burst structure declined in parallel. Before muscimol, multi-peak bursts retained the characteristic cascading profile. After muscimol, this profile was progressively lost, high-peak-count bursts became less frequent, intra-burst peak amplitudes decreased and multi-peak event rates declined across the post-injection period (Fig. 2l–p). Thus, muscimol produced a graded reduction in both the LFP and MUA expression of pacemaking before population bursts disappeared.

### Disinhibition produces a distinct epileptiform regime

The physiological model described the effects of reduced inhibition. According to the model, blocking GABA-A receptor signaling under baseline conditions should not produce the same pacemaker regime. Glutamate antagonism reduces ongoing recruitment and permits recovery-dependent initiation; disinhibition preserves glutamatergic excitation while removing inhibitory restraint. The expected model output is therefore epileptiform activity, not the same recovery-paced pacemaker.

Picrotoxin produced this distinct state. Simulations with reduced GABAergic conductance generated rapid, synchronized, non-cyclic population activity (Fig. 3a,b). In vivo, local picrotoxin under baseline conditions produced broad disinhibited bursts in local-field-potential and multi-unit recordings (Fig. 3c,d). Compared with DGG, picrotoxin events had broader burst-duration and suppression-duration distributions and more prolonged spike trains within bursts (Fig. 3e-g). The mean intra-burst amplitude profile also differed: rather than the declining pacemaker cascade, picrotoxin events rose toward a mid-event maximum and then declined (Fig. 3h).

**Figure 3.**
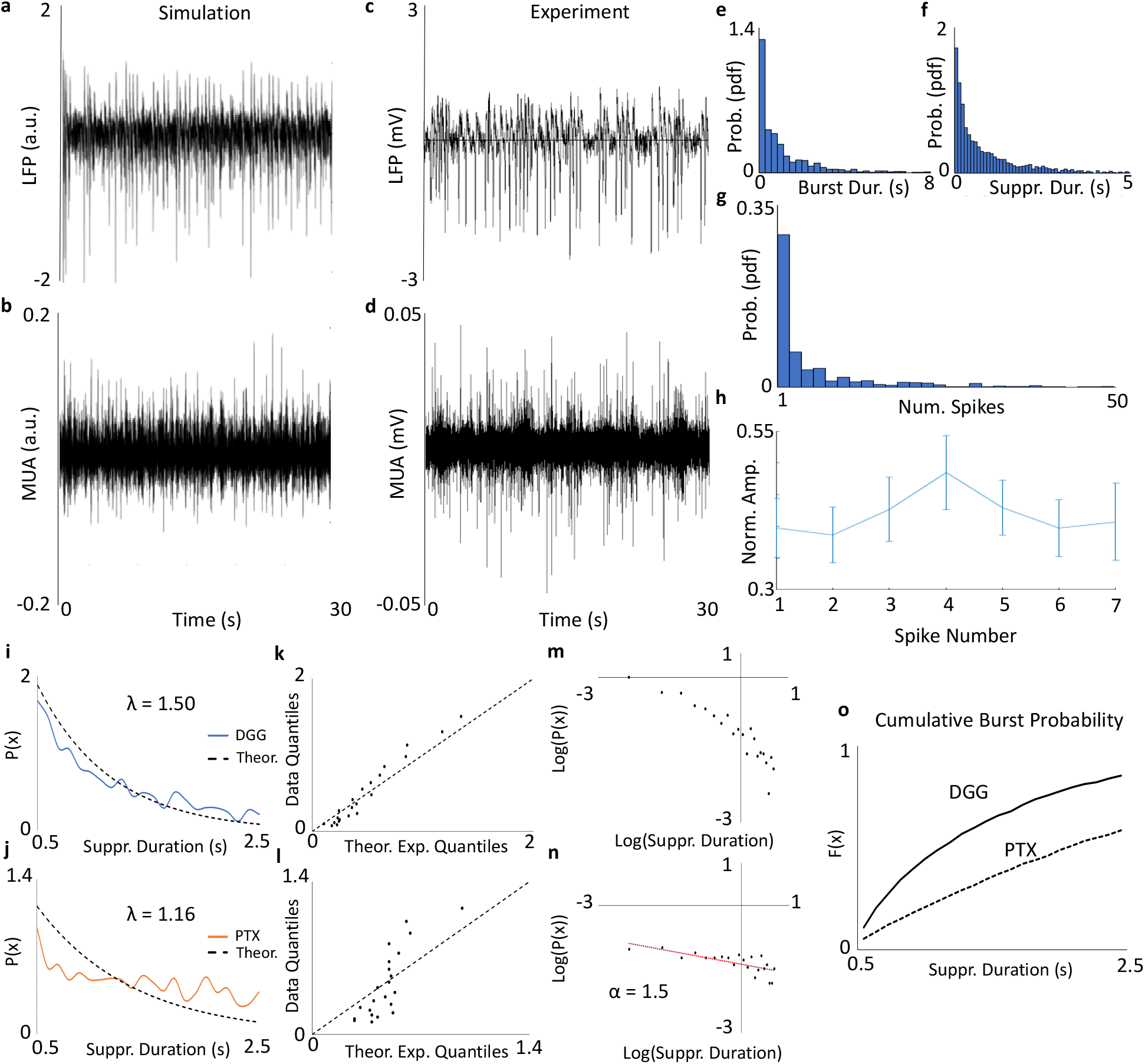
Picrotoxin produces epileptiform activity, not pacemaking. a,b, Simulations with reduced GABAergic conductance produce rapid, non-cyclic population activity. c,d, Picrotoxin recordings in vivo show the corresponding disinhibited state in LFP and MUA. e-g, Picrotoxin produces broad distributions of burst duration, suppression duration and intra-burst spike number, indicating more variable and prolonged recruitment than the pacemaker regime. h, The mean intra-burst amplitude profile rises toward a mid-burst maximum and then declines, contrasting with the declining pacemaker cascade. i,j, Shifted exponential fits to truncated suppression-duration distributions for DGG and picrotoxin over the common 0.5-2.5 s support. k,l, Exponential quantile plots for DGG and picrotoxin. m,n, Log-log plots of suppression-duration structure; picrotoxin shows heavier-tailed, power-law-like scaling (alpha = 1.5). o, Cumulative next-burst probability as a function of time since the previous burst. These analyses distinguish the DGG post-recovery waiting-time structure over the analysed range from disinhibited picrotoxin recurrence.

The waiting-time analysis strengthened this distinction. For a direct comparison, suppression durations were analyzed on the same truncated range of 0.5-2.5 s and shifted so that the 0.5 s suppression cutoff corresponded to zero. On this support, DGG was closer to a shifted exponential waiting process (lambda = 1.50), whereas picrotoxin showed weaker exponential agreement (lambda = 1.16; Fig. 3i-l). Log-probability and log-log analyses further separated the regimes: DGG did not show strong linear scaling in log-log space, whereas picrotoxin showed heavier-tailed, power-law-like scaling (alpha = 1.5; Fig. 3m,n). The cumulative next-burst probability curves also differed across time since the previous burst (Fig. 3o).

Together, the pharmacological manipulations produced three distinct population patterns. Glutamate-receptor antagonism produced recurrent burst–quiescence, muscimol progressively reduced and ultimately abolished those bursts, and picrotoxin produced broad disinhibited epileptiform activity with different timing and intra-burst structure. The formal availability-recruitment model is provided in Supplementary Methods 4.

## Discussion

The main finding is that a reservoir hypothesis derived from two previously unconnected single-neuron constraints predicted a hidden cortical operating regime that was then observed experimentally. To our knowledge, this is the first demonstration that local ionotropic glutamate-receptor antagonism alone can induce sustained burst-quiescence in awake cortex without preceding disinhibition or seizure induction. Glutamate-receptor antagonism revealed recovery-dependent pacemaking, muscimol progressively reduced its recurrence and amplitude before suppressing it, and picrotoxin produced a quantitatively distinct epileptiform regime. These are not three unrelated drug effects; they define complementary paced, quiescent and epileptiform operating states of the same class of cortical tissue. In this sense, the cortex contains the components of a population pacemaker machine, but ongoing glutamatergic activity normally prevents those components from operating together.

The model does not require a unique molecular or anatomical trigger for each burst. Residual afferent, neuromodulatory, intrinsic or extracellular fluctuations may provide the immediate perturbation that initiates an event. Crucially, such inputs need not be rhythmic: they generate a population burst only when the local cortical population has recovered sufficiently and recruitment can cross threshold. The variable waiting-time structure, local intra-burst LFP cascade and muscimol-dependent changes in both burst size and recurrence support state-dependent cortical generation.

We call the broader mechanism the excitation-interruption principle. Excitation does not only drive activity; when continuous, it can suppress population timing by repeatedly recruiting spikes and resetting recovery. Reducing excitation can therefore expose rhythmic population output rather than eliminate it. Muscimol progressively reduced burst amplitude, MUA recruitment structure and recurrence before abolishing the pacemaker regime, whereas disinhibition produced epileptiform activity.

The muscimol transition provided a direct operational test of the availability reservoir. LFP reports summed local synaptic and transmembrane activity, whereas MUA reports action potentials from the recruited local population. Their parallel decline showed that a progressively smaller fraction of local population activity was recruited into each burst. Burst recurrence fell and the intervals between successful bursts lengthened at the same time, showing that the smaller reservoir supported successful initiation less often. The disappearance of both LFP and MUA bursts marked the lower functional limit at which the reservoir could no longer express pacemaking. These signals measure reservoir participation relative to established pre-muscimol pacemaking rather than an absolute number of neurons.

These findings recast pacemaking as a population-state property rather than an exclusive property of specialized cell types. Existing uses of the term “pacemaking” span intrinsically bursting neurons, rhythmogenic circuits, coupled pacemaker tissues and self-organized firing clusters in excitable media.^3,9–11^ Intrinsic rhythmicity of single cells is insufficient to explain coherent population timing; coherent output additionally requires synchronization and population recruitment.^2,12^ The availability reservoir supplies this population-level coordination: each event transiently depletes eligibility, recovery rebuilds it and thresholded recruitment converts the recovered pool into the next event.

This framework also clarifies why the electrophysiological pattern alone cannot identify mechanism. The pacemaker regime can resemble conventional anesthesia-induced burst suppression, and disinhibited epileptiform activity can likewise appear as large bursts separated by quiescent intervals. ^13–15^ However, all recordings in the present study were obtained in fully awake animals; anesthesia was therefore not a component of the experimental state. We consequently used picrotoxin-induced disinhibition as the relevant awake-cortex comparator.

Quantitative comparison showed that superficially similar burst–quiescence states were distinguished by pharmacological direction, waiting-time and event-duration distributions, and intra-burst recruitment physiology (Figs. 1 and 3). Thus, burst–quiescence is not a single mechanistic state: waveform appearance alone cannot distinguish disinhibited epileptiform activity and recovery-dependent pacemaking unless the underlying dynamics are analyzed.

Conventional acute-slice paradigms are also not equivalent to the intact awake preparation studied here. The proposed mechanism may depend on a sufficiently large, intact three-dimensional population.

The same control logic may extend beyond the cortex. Canonical pacemakers are often described by specialized cells or circuits, yet stable population rhythm requires more than cellular automaticity: after each event, participating tissue must become temporarily unavailable, and the next event must wait for recovery. Published heart and respiratory recovery measurements can be interpreted on the same availability-recruitment axes.^16,17^ The cortical result extends this logic to a tissue not usually treated as pacemaking, suggesting that recovery-gated recruitment can be a general organizing motif of excitable systems without requiring biophysical identity across organs.

The study establishes a previously unrecognized type of cortical population activity in the awake brain. The regime was absent at baseline, consisted of structured compound bursts separated by quiescence, was progressively titrated by GABAergic inhibition and was quantitatively distinct from disinhibition-induced epileptiform activity. We identify this operating state as recovery-dependent cortical population pacemaking. Future work should test how broadly this principle applies across brain regions, cardiac tissues, species and physiological or pathological states, and define the minimum population size, tissue geometry and recruitment strength required for its emergence.

## Methods

### Animals and experimental design

Adult Dutch-belted rabbits (2.5–3.5 kg; 8–12 months of age) were studied in accordance with National Institutes of Health guidelines and protocols approved by the Endeavor Health (formerly NorthShore University HealthSystem) Institutional Animal Care and Use Committee. Three pharmacological groups were used (n = 6 per group). Group size was based on prior electrophysiological studies using the same chronic rabbit somatosensory-cortex preparation rather than on a formal prospective power calculation.^6–8^

In the DGG-only group, a 15-min awake baseline recording was followed by local gamma-D-glutamylglycine (DGG) injection, and recording continued until 50 min from recording onset. In the DGG–muscimol group, DGG was administered after the same 15-min baseline. Muscimol was administered only after the DGG-induced pacemaker regime had remained stable for at least 10 min. Because the time required for stabilization varied among recordings, the absolute time of muscimol administration varied among animals; all muscimol analyses were aligned to the time of muscimol injection, and recording continued for at least 20 min thereafter. In the picrotoxin group, picrotoxin was administered after a 15-min baseline and recording continued until 50 min from recording onset.

### Surgery and chronic electrophysiology

Animals were anesthetized for surgery with subcutaneous ketamine (60 mg kg−1) and xylazine (10 mg kg−1). Four platinum–iridium microwires (25 µm diameter, heavy-Formvar coated; California Fine Wire) and a 250-µm silica injection cannula (Polymicro Technologies) were chronically implanted into layers IV–V of somatosensory cortex using previously reported coordinates and procedures.^6^ The electrode–cannula assembly was mounted on a custom nylon microdrive that permitted vertical adjustment of electrode position.^7^ A 150-µm silver wire positioned between skull and dura served as the reference electrode. Animals recovered for at least 1 week before habituation and recording.

Recordings were obtained in awake animals after habituation to light restraint in cloth bags. Animals were not head-fixed. Microwire signals were amplified through a Neuralynx miniature preamplifier and multichannel differential amplifier system (Neuralynx). Spike channels were band-pass filtered at 0.3–3 kHz, local-field-potential (LFP) channels were filtered at 1–200 Hz, and all signals were digitized at 32 kHz per channel. LFP and multi-unit activity (MUA) were recorded continuously. Before drug administration, the primary analysis electrode was selected as the electrode with the largest stable baseline MUA signal and the closest effective proximity to the injection site.

### Electrode localization

The chronic electrode and cannula positions were verified by magnetic resonance imaging in awake animals using a 9.4-T BioSpec 94/30USR imaging spectrometer (Bruker) operating at a proton frequency of 400 MHz. A 20-mm circular surface coil was used for radiofrequency transmission and reception. Anatomical images were acquired with a multislice gradient-echo sequence using a 512 × 512 matrix, a 48 × 48 mm field of view (94 × 94 µm in-plane resolution), 1.0-mm slice thickness, repetition time of 1.5 s, echo time of 20 ms and eight signal averages, following the established imaging protocol.^6,7^

### Local pharmacology

DGG (gamma-D-glutamylglycine; Tocris) was used as a broad ionotropic glutamate-receptor antagonist. DGG was prepared at 0.098 µmol µl−1 and 1 µl was injected intracortically into somatosensory cortex after the 15-min baseline. The concentration was selected based on testing various concentrations and from prior work demonstrating strong blockade of ionotropic glutamatergic transmission at this concentration.^18^ We also tested the selective NMDA antagonist 2-amino-5-phosphonopentanoic acid (AP5, 12.68 mM), which produced a similar effect to DGG but with a shorter duration (up to 10-15 minutes). Due to its longer-lasting effect, DGG was chosen for these experiments. Muscimol was used to increase GABA_A-receptor signaling after establishment of the pacemaker regime. Muscimol (0.4 g l−1; 1 µl) was injected only after at least 10 continuous minutes of stable DGG-induced burst–quiescence. The 10 min immediately preceding muscimol constituted the pre-muscimol reference period. Post-muscimol analyses were expressed relative to the injection time rather than to absolute recording time.

Picrotoxin (248 µM) was dissolved in saline and administered under baseline conditions to produce local disinhibition while avoiding generalized convulsive seizures. All injections were 1 µl and were delivered through a silica tube or needle (190 µm outer diameter, 100 µm inner diameter; Polymicro Technologies) connected through transparent Tygon tubing to a Hamilton syringe. Equal-volume saline vehicle injections were performed in the same preparation using the same procedure, and multi-unit activity was monitored to minimize injection-volume artefacts.^7^ Previous work using this preparation and injection volume showed that vehicle injection alone did not alter LFP or MUA activity.^6–8^

### MUA extraction

MUA event extraction used the standard Neuralynx workflow and the same pipeline for all experimental conditions (baseline, DGG, DGG plus muscimol, PTX and saline vehicle). MUA bursts were detected by threshold crossing in Neuralynx. ^8^ The resulting MUA bursts were used separately to quantify firing and intra-burst recruitment within LFP-defined events, while the burst-quiescent periods were defined from LFP recordings.

### Burst and suppression segmentation

Burst and suppression periods were segmented from normalized LFP traces using the same operational definitions across DGG and picrotoxin conditions. Each LFP trace was converted to z scores. If the LFP trace stays within a 2 standard deviation threshold for at least 500 ms, then the trace is defined as in a suppression phase, which lasts until the trace crosses the two-standard deviation threshold. The traces which are in between the suppression periods are candidates for the burst phase. If these candidate phases are at least 250 ms and under 30 seconds, then it was considered a burst phase. The 500-ms minimum separated suppression periods from short gaps within compound bursts and was applied identically across conditions.

For stable DGG and picrotoxin analyses, events were evaluated from 5 to 25 min after injection (20–40 min from recording onset). For every burst, the analysis stored onset and offset, duration, LFP amplitude, number of intra-burst LFP peaks, and intra-burst firing frequency. Mean LFP intra-burst amplitude profiles were evaluated for four-peak pacemaking bursts and seven-peak picrotoxin bursts, corresponding to the characteristic peak counts used in the respective experimental summaries.

### Muscimol analysis

Muscimol analyses were aligned to injection because the injection time depended on the establishment of at least 10 min of stable pacemaking. The pre-muscimol reference comprised the final 10 min of stable pacemaking immediately before injection. Representative LFP traces were selected from 1 min before muscimol and from 5 and 10 min after injection. Post-muscimol intra-burst amplitude profiles were summarized in successive 2-min windows, whereas peak-count proportions and multi-peak event rates were summarized in successive 5-min windows through 20 min after injection. LFP burst amplitude, burst recurrence and LFP intra-burst structure was normalized to the stable pre-muscimol period where indicated.

### Statistical Analyses

To analyze the statistical properties of pacemaking bursts, the density distributions were plotted for intra-burst duration, suppression duration, and the number of intra-burst spikes. The fit of the suppression durations was checked with the theoretical exponential probability density function (pdf) using maximum likelihood estimation (MLE), since suppression times follow the logic of waiting times. Suppression durations greater than 10 seconds were excluded for the DGG analysis. Both duration and the number of intra-bust spikes however were expected to follow a log-normal distribution since both metrics typically have a strong right skew and both are strictly positive values. Both the mu and the sigma parameters were estimated using the mean and standard deviation of the log of the metric data. The average burst rate (Hz) was calculated for each of the 6 experiments, and the distribution via a box-whisker plot was included.

Every burst of every spike amount was recorded across all 18 experiments (DGG, MSC, and PTX). For DGG, for bursts that had only four pronounced spikes, we recorded the average amplitude per each spike and then used a paired sample t-test to confirm the significance of the difference in the amplitude for each spike number. Similarly for each experiment, the average burst duration, average post suppression initiation rate (PSIR), average burst amplitude, and average suppression duration were calculated from minute 17 to minute 18 (shortly after DGG injection) and from minute 24-25 (ten minutes after DGG injection), and a paired two-tailed t-test was run to determine significant differences in those metrics. For each one-minute time window starting from 17 minutes of the recording up until 25 minutes of the recording, the lambda parameter of the fitted exponential distribution on suppression durations, the mu and sigma parameters for the fitted log-normal distribution for the burst duration metrics, and the mu parameters for the fitted log-normal distributions for the number of intra-burst spikes were calculated and plotted over time.

For the MSC experiments, we ran statistics on two main parameters: The relative change in amplitude relative to a pre-injection baseline and the change in PSIR (Hz). A two-sample t-test was run to compare the relative baseline amplitude of the control (DGG only) against the treatment (DGG + MSC), while a paired t-test was run to analyze the change in the PSIR before and after injection of the MSC group. The timing for comparison of burst amplitudes between the DGG and the DGG+MSC groups started at 25 minutes for the DGG group and the timing of the MSC injection, for each group respectively. To compare the changes between the PSIR before and after injection of MSC in the DGG experiments, we ran a paired two-tailed t-test.

### Conceptual availability–recruitment model

The conceptual model separated recovery of availability from initiation of the next population event. The availability variable represented the relative amount of local tissue that was simultaneously susceptible and sufficiently recovered. Between bursts, availability recovered toward its upper bound with an effective recovery time constant; each burst subsequently depleted availability by an event-size term. Once a prespecified recovery condition was reached, the exact event time was determined by a stochastic initiation component. Full equations and implementation details are provided in Supplementary Methods 4.

The pharmacological conditions were mapped onto this reduced framework qualitatively. DGG reduced ongoing glutamatergic recruitment and permitted availability to rebuild. Muscimol increased inhibitory restraint and progressively reduced expression of the recovered population as a propagating burst. Picrotoxin reduced inhibitory restraint and shifted the tissue toward disinhibited epileptiform recruitment. It was represented in the physiological model and used experimentally as a distinct comparator.

### Physiological cortical model

Physiological simulations adapted a previously described two-compartment soma–dendrite model.^19,20^ Somatic and dendritic membrane potentials were governed by Hodgkin–Huxley-type currents, passive soma–dendrite coupling, synaptic conductances and extracellularly induced transmembrane perturbations. The soma contained transient sodium and delayed-rectifier potassium currents. The dendrite contained persistent sodium and potassium currents that supported heterogeneous subthreshold susceptibility oscillations.^21^ Synaptic terms included recurrent and external glutamate-mediated excitation, tonic and phasic GABAergic conductances and spike-history-dependent recovery. The complete membrane equations, gating functions, conductance parameters and equilibrium potentials are provided in Supplementary Methods 5 and Supplementary Tables 1 and 2.

The simulated intact cortical volume was 1,000 × 250 × 500 µm and was represented by a 20 × 2 × 10 grid. The x–z plane represented 200 model neurons and the two positions along the y axis represented soma and dendrite compartments. Somatic injected current was set to −0.45 µA cm^−2^ with synaptic noise and a neuron-specific recovery current dependent on the time since the most recent action potential; dendritic injected current was set to 0.8 µA cm^−2^ to support subthreshold oscillations.^5,21^ The coupled equations were integrated by the forward-Euler method with a 0.1-ms timestep.

Extracellular potential at the virtual recording site was calculated as a distance-weighted sum of somatic and dendritic transmembrane currents, following established field-potential and ephaptic-coupling approaches.^22–24^ Simulations with and without extracellular-potential feedback were compared to test its contribution to synchrony and compound-burst amplitude. This comparison evaluated one plausible recruitment route and did not make extracellular coupling a required component of the conceptual reservoir model.

Resting, DGG, DGG–muscimol and picrotoxin protocols were generated from the same model architecture. DGG was simulated by removing glutamate-mediated recruitment while retaining the remaining resting-state parameters. In the sequential DGG–muscimol protocol, muscimol was introduced only after recurrent DGG pacemaking was established, and tonic GABAergic conductance was then increased over the specified transition period. Picrotoxin was simulated by reducing GABA_A_-mediated and tonic inhibitory restraint while retaining glutamatergic recruitment. Detailed parameter values, intervention timing, randomization and signal-construction procedures are provided in Supplementary Methods 5 and 6 and in the accompanying code.

## Data availability

Data will be deposited in a public repository before publication.

## Code availability

The physiological-model code will be deposited in a public repository before publication.

## Use of language models

Generative language-model assistance was used during manuscript preparation for language editing and review. All scientific content, analyses, code, interpretations and final wording were reviewed and approved by the authors, who take full responsibility for the work.

## Author Contributions

D.P.A. conceived the study, developed the conceptual framework, designed and supervised the experiments, interpreted the data and wrote the manuscript. E.D. performed the quantitative analyses, developed and implemented the computational models, prepared figures and contributed to manuscript preparation. N.A.S. contributed to the experimental work and data acquisition and critical revision of the manuscript. M.M.M. and S.B.G. contributed to interpretation of the findings and critical revision of the manuscript. All authors reviewed and approved the final manuscript.

## Additional Information

Supplementary Information is available for this paper.

