## Supplemental information for "Glutamatergic excitation suppresses latent cortical pacemaking"

### Contents

Supplementary Methods 1. Animal preparation and recording
Supplementary Methods 2. Pharmacology and injection controls
Supplementary Methods 3. Event detection and multi-level statistics
Supplementary Methods 4. Conceptual availability-reservoir model
Supplementary Methods 5. Physiological soma-dendrite model
Supplementary Methods 6. Network and extracellular-potential model
Supplementary Tables 1-2. Model parameters
Supplementary Note 1. Truncated waiting-time comparison between DGG and picrotoxin

Supplementary References

### Supplementary Methods 1. Animal preparation and recording

Adult Dutch-belted rabbits (2.5–3.5 kg, 8–12 months of age) were used. All procedures conformed to National Institutes of Health guidelines and were approved by the Endeavor Health (formerly NorthShore University HealthSystem) Institutional Animal Care and Use Committee.

Three groups of animals were used. In the first group (N = 6), DGG was injected after a 15-min baseline recording, and electrophysiological recording continued until 50 min from recording onset. In the second group (N = 6), DGG was injected after the same baseline period, and muscimol (MSC) was administered only after DGG-induced pacemaking had remained stable for at least 10 continuous minutes. Because stabilization time varied among recordings, all muscimol analyses were aligned to the time of MSC injection, and recording continued for at least 20 min thereafter. In the third group (N = 6), picrotoxin (PTX) was injected after a 15-min baseline recording, and recording continued until 50 min from recording onset.

The animals were anesthetized with a subcutaneous injection of ketamine (60 mg/kg) and xylazine (10 mg/kg) and underwent stereotaxic surgery for chronic electrode and injection-cannula implantation into somatosensory cortex, as described previously.[1] Four platinum–iridium electrodes (25 μm each, coated with heavy Formvar; California Fine Wire, Grover Beach, CA) and a 250-μm silica injection cannula (Polymicro Technologies, Phoenix, AZ) were implanted into layers IV–V of somatosensory cortex using previously reported coordinates.[1] The assembly was attached to a custom-made nylon microdrive that permitted vertical adjustment of electrode position.[2] A 150-μm silver wire was placed between the skull and dura mater to serve as a reference electrode. After surgery, the animals recovered for at least one week before habituation and recording.

Electrophysiological signals were recorded chronically in awake animals lightly restrained in cloth bags without head fixation after habituation to the recording setup. Signals from the microwires were amplified through a miniature preamplifier and multichannel differential amplifier system (Neuralynx, Inc., Bozeman, MT, USA), band-pass filtered at 0.3–3 kHz for spike channels and 1–200 Hz for local field potentials, and digitized at 32 kHz per channel. Multi-unit activity (MUA) and local field potentials (LFP) were recorded continuously. The primary electrode was prespecified as the electrode with the largest stable baseline MUA signal.

### Supplementary Methods 2. Pharmacology and injection controls

Based on prior studies using the same chronic rabbit preparation, a sample size of N = 6 rabbits per group was used.[1–3] Muscimol has been used extensively for local GABA-A-receptor activation in vivo.[1,4,5]

Experiments combining DGG and muscimol (MSC) were conducted (N = 6) to test whether increasing GABA-A-receptor signalling suppresses established pacemaking. MSC was administered at 0.4 g/l only after DGG-induced pacemaking had remained stable for at least 10 continuous minutes; the absolute injection time therefore varied among recordings.

DGG (gamma-D-glutamylglycine; Tocris, Minneapolis, MN, USA; 1 µl) was used to block ionotropic glutamatergic transmission at AMPA and NMDA receptors. A concentration of 0.098 µmol/µl was selected from prior work demonstrating strong glutamatergic blockade at this dose.[6] Following a 15-min baseline recording, DGG was injected intracortically into somatosensory cortex. Muscimol (0.4 g/l; 1 µl) was administered after at least 10 continuous minutes of stable DGG-induced pacemaking to test whether burst–quiescence could be suppressed through GABA-A-receptor activation. All muscimol analyses were aligned to the time of injection.

We also tested the selective NMDA-receptor antagonist 2-amino-5-phosphonopentanoic acid (AP5; 12.68 mM), which produced a qualitatively similar but shorter-lived effect than DGG (up to 15 min). DGG was selected for the principal experiments because its effect was longer lasting.

Picrotoxin (PTX; 248 µM) was dissolved in saline. This concentration was selected to avoid generalized convulsive seizures while allowing local disinhibition.[3,7] All injections (1 µl) were delivered through a silica tube or needle (190 µm outer diameter and 100 µm inner diameter; Polymicro Technologies, LLC, Phoenix, AZ) connected to a Hamilton syringe through transparent Tygon tubing. Equal volumes of saline vehicle were injected using the same procedure to control for injection-related effects. Multi-units were monitored to minimize volume effects.[2] Neuronal activity was recorded before, during and after pharmacological modulation. Previous work and the DGG experiments indicated that 15 min was sufficient to avoid injection-volume artefacts and allow local diffusion.[2]

### Supplementary Methods 3. Event detection and multi-level statistics

MUA event extraction used the standard Neuralynx workflow and the same pipeline for all experimental conditions (baseline, DGG, DGG plus muscimol, PTX and saline vehicle). MUA bursts were detected by threshold crossing in Neuralynx.[3] The resulting MUA bursts were used to quantify firing and intra-burst recruitment within LFP-defined events.

Statistical analyses were performed at multiple levels because the questions differed across levels. Drug effects across conditions were evaluated with the animal or injection as the biological experimental unit. Event-generation mechanisms were evaluated at the burst level because burst duration, suppression duration, intra-burst peak count and waiting-time distributions distinguish DGG pacemaking from PTX disinhibition. Burst-level statistics were therefore reported as event-level evidence. For PTX, the suppression-duration power-law exponent was estimated by maximum likelihood, and log–log plots were used to visualize scaling and compare PTX with DGG.

To distinguish suppression periods from short gaps within compound bursts, the minimum suppression period was set to 500 ms. After each LFP trace was converted to z scores, a suppression period was defined as at least 500 ms of activity within the 2-standard-deviation threshold during the interval from 5 to 25 min after injection (20–40 min from recording onset). Suppression duration was measured from the resulting suppression vectors. For time intervals where LFP activity was not in a suppression phase, if the interval lasted longer than 250 ms that it was classified as a bursting period; bursts longer than 30 s were excluded. Each burst vector was retained to calculate burst duration, number of intra-burst MUA peaks and burst firing frequency. Events were further classified by peak number to calculate mean intra-burst peak amplitudes. For DGG, intra-burst amplitudes were evaluated in four-peak bursts, and for PTX they were evaluated in seven-peak bursts, corresponding to the characteristic peak counts used in the respective experimental summaries. Paired t-tests were used for the stated intra-experiment burst- and suppression-parameter comparisons, with the exceptions of intra-experiment comparisons .

Electrode localization was performed using MRI (Figure S1). Imaging was performed in awake animals using a 9.4-T BioSpec 94/30USR imaging spectrometer (Bruker, Billerica, MA, USA) operating at a proton frequency of 400 MHz. A flat circular surface coil (20 mm diameter) was used for radiofrequency transmission and reception. High-resolution anatomical images were obtained using a multislice gradient-echo sequence (512 × 512 matrix, 48 × 48 mm field of view, 94 × 94 µm in-plane resolution, 1.0-mm slice thickness, TR 1.5 s, TE 20 ms and eight signal averages) following the established imaging approach.[1,2]

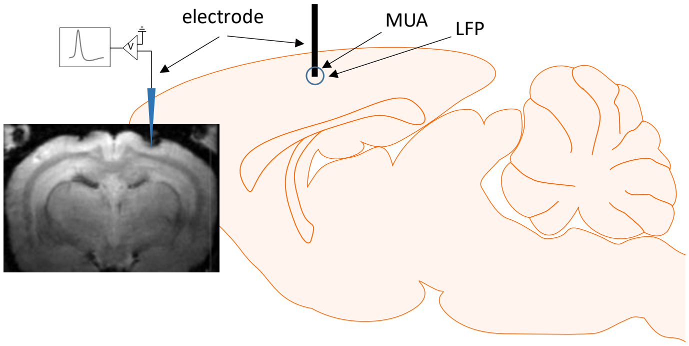


*Figure S1. The concept of the recording. The electrodes were implanted into somatosensory cortex, as illustrated by the MR image, to record MUA and LFP.*

### Supplementary Methods 4. Conceptual availability-reservoir model

The conceptual model separates recovery of availability from initiation of the next population event. Let $A_k$ denote the effective availability reservoir, the fraction of local tissue that is both susceptible and sufficiently recovered at time bin k. Between bursts, availability recovers toward one with recovery time constant $\tau_{rec}$; a burst depletes availability by an amount $b_k$.

$$A_{k+1} = A_k + \Delta t\left(\frac{1-A_k}{\tau_{\mathrm{rec}}}\right)$$

We define, a priori, the availability threshold $0 < \theta < 1$, where once $A_k > \theta$ the chance of a burst follows the dynamics of $\varphi_k$ (starting at $\varphi_k = 0$):

$$\varphi_{k+1} = \varphi_k + \Delta\mathrm{t}(\lambda(1-\varphi_k))$$

Where $\lambda$ denotes the rate parameter controlling the average waiting time for a burst occurrence. At every time step k, a value is selected uniformly from [0,1]. If $\varphi_k$ exceeds the selected value, the population bursts and $\varphi_k$ is reset to zero. This phenomenological stochastic component generates variable post-recovery event times; experimental waiting-time distributions were fitted separately. The burst-size term $b_k$ was selected within the permitted interval from $\theta$ to $A_k$, after which availability was updated as $A_{k+1} := A_k - b_k$.

Thus, $b_k$ is not the trigger for burst initiation; it is the depletion caused by a burst after initiation. In a deterministic alternative, a burst occurs when $A_k$ crosses a fixed recruitment threshold, predicting that larger bursts should be followed by longer suppression intervals. In the current formulation, recovery creates the permissive state, but the exact burst time is drawn from a probabilistic process. For the DGG data, suppression durations after the 500-ms operational cutoff were fitted separately with an exponential waiting-time distribution.

In pharmacological terms, DGG reduces ongoing glutamatergic recruitment and allows $A_k$ to rebuild; MSC increases GABA-A-mediated inhibition and lowers effective biological recruitment below the propagation threshold. The reduced conceptual model formalizes recovery-dependent pacemaking and its progressive loss under increased inhibition. PTX-induced disinhibited activity is represented only in the physiological model and is used experimentally as a distinct comparator.

### Supplementary Methods 5. Physiological soma-dendrite model

The computational neuronal firing patterns were calculated by adapting a previously established soma–dendrite model.[7,8] The transmembrane potential was governed by the equations

$$\frac{C_s(dV_{m,s} + dV_{e,s})}{dt} = -I_{ion,s} - I_{syn,s} - I_{res,s} + I_{inj,s} + I_{ds}$$

$$\frac{C_d\left(dV_{m,d} + dV_{e,d}\right)}{dt} = -I_{ion,d} - I_{syn,d} - I_{res,d} + I_{inj,d} - I_{ds}$$

where $V_{m,s}$ and $V_{m,d}$ (in mV) are the transmembrane potentials for the soma and dendrite, respectively, and $dV_{e,s}$ and $dV_{e,d}$ are perturbative terms (for the soma and dendrite compartments, respectively), to account for the transmembrane capacitive current induced by extra-cellular potential fluctuations.[9,10] The injected current into the soma and dendrite, respectively, is denoted by $I_{inj,s}$ and $I_{inj,d}$. The resistive current due to extra-cellular potential fluctuations is denoted by $I_{res,s}$ and $I_{res,d}$, for the soma and dendrite compartments, respectively.

The capacitance for the soma and dendrite compartment are represented by $C_s = 3$ and $C_d = 0.25$ (in μF/cm²) The current between the passive current between the soma-dendrite compartments is represented by $I_{ds}$, and is defined as $I_{ds} \coloneqq g_c(V_{m,d} - V_{m,s})$, where $g_c$ (in mS/cm2) is the coupling strength between the soma and dendrite compartments.[8] In accordance with,[11] the equation for the resistive extra-cellular current is given by:

$$I_{res,s} = \left(\overline{g}_K + \overline{g}_{Na} + \overline{g}_{leak,s}\right) V_{e,s}$$

$$I_{res,d} = \left(\overline{g}_{K-P} + \overline{g}_{Na-P} + \overline{g}_{leak,d}\right) V_{e,d}$$

The soma compartment contained an inward sodium current and an outward delayed-rectifier potassium current,[8] whereas the dendritic compartment was driven primarily by weak persistent sodium and potassium currents supporting subthreshold oscillations.[12]

The ion channel current for the soma, $I_{ion,s}$, is determined by:

$$I_{ion,s} = I_K + I_{Na} + I_{leak,s}$$

Where

$$I_K = \overline{g}_K n\left(V_{m,s} - E_K\right)$$

$$I_{Na} = \overline{g}_{Na} m_\infty^2 h\left(V_{m,s} - E_{Na}\right)$$

$$I_{leak,s} = g_{leak,s}\left(V_{m,s} - E_{leak}\right)$$

And the ion channel current for the dendrite, $I_{ion,d}$, is determined by:

$$I_{ion,d} = I_{K-P} + I_{Na-P} + I_{Ca} + I_{leak,d}$$

Where

$$I_{KP} = \overline{g}_{KP} n_p\left(V_{m,d} - E_K\right)$$

$$I_{NaP} = \overline{g}_{NaP} m_p\left(V_{m,d} - E_{Na}\right)$$

$$I_{Ca} = \overline{g}_{Ca} \alpha\left(V_{m,d} - E_{Ca}\right)$$

$$I_{leak,d} = g_{leak,d}\left(V_{m,d} - E_{leak}\right)$$

The kinetic equations for each gating variable are given by

$$\frac{dn}{dt} = \left(n_\infty\left(V_{m,s}\right) - n\right)/\tau\left(V_{m,s}\right)$$

$$\frac{dh}{dt} = \left(h_\infty\left(V_{m,s}\right) - h\right)/\tau\left(V_{m,s}\right)$$

$$\frac{dn_p}{dt} = \left(n_{p,\infty}\left(V_{m,d}\right) - n_p\right)/\tau_{np}\left(V_{m,d}, T\right)$$

$$\frac{dm_p}{dt} = \left(m_{p,\infty}\left(V_{m,d}\right) - m_p\right)/\tau_{mp}$$

The steady state values for each gating variable are in the form $y_\infty = \alpha_y/\left(\alpha_y + \beta_y\right)$, while the time constants are in the form $\tau = 1/\left(\alpha_y + \beta_y\right)$ for $y = h, n, m$. The functions $\alpha_y$ and $\beta_y$ for all values of $y$ are

$$\alpha_h = 0.128 \exp\left(\frac{17 - V_{m,s}}{18}\right)$$

$$\beta_h = \frac{4}{\exp\left(\frac{40 - V_{m,s}}{5}\right) + 1}$$

$$\alpha_m = \frac{0.32\left(13.1 - V_{m,s}\right)}{\exp\left(\frac{13.1 - V_{m,s}}{4}\right) - 1}$$

$$\beta_m = \frac{0.28\left(V_{m,s} - 40.1\right)}{\exp\left(\frac{V_{m,s} - 40.1}{5}\right) - 1}$$

$$\alpha_n = \frac{0.016\left(35.1 - V_{m,s}\right)}{\exp\left(\frac{35.1 - V_{m,s}}{5}\right) - 1}$$

$$\beta_n = 0.25 \exp\left(0.5 - 0.025 V_{m,s}\right)$$

In accordance with previously described persistent sodium and potassium mechanisms underlying subthreshold oscillations,[12]

$$n_{p,\infty} = \frac{1}{(1 + \exp\left(-(V_{m,d} + 35)/10\right)}$$

$$m_{p,\infty} = \frac{1}{(1 + \exp\left(-(V_{m,d} + 40)/5\right)}$$

$$\tau_{np} = \frac{1000}{3.3 \exp\left(\frac{V_{m,d} + 35}{40}\right) + \exp\left(\frac{-\left(V_{m,d} + 35\right)}{20}\right)\left(\frac{1}{3\hat{}((T - \mu - 22)/10)}\right)}$$

$$\tau_{mp} = 1$$

Where T = 32 °C is the temperature of the medium and $\mu$ is a scaling value uniformly distributed between [0,10] assigned to each neuron to produce subthreshold oscillations between 4–12 Hz.

The calcium-spike gating variable was modelled as a rise-and-decay function after a somatic action potential through back-propagation.[13]

$$\alpha = \frac{\exp(1)(t-t')}{10} exp\left(-\frac{(t-t')}{10}\right)$$

Where $t'$ is the time of the most recent somatic action potential.

Both compartments contain $I_{GABA}$ as a GABA-mediated current. The leak current for both compartments is denoted by $I_{leak}$. All currents are in units of μA/cm$^2$.

The equations of the synaptic currents for the soma, $I_{syn,s}$, are:

$$I_{syn,s} = I_{GABA-A} + I_{Tonic} + I_{Glu}$$

Where

$$I_{GABA-A} = g_{GABA-A}(V_{m,s} - E_{Cl})$$

$$I_{GABA-Ton} = g_{Tonic}(V_{m,s} - E_{GABA})$$

$$I_{Glu,S} = g_{Glu}(V_{m,s} - E_{Glu})$$

Tonic GABA dynamics were based on established descriptions of population GABA signaling and tonic GABA-A conductance.[14,15]

$$\frac{dg_{Tonic}}{dt} = \gamma(f_m(t) + f_b) - \omega(g_{Tonic} - g_{basal})$$

Where $\gamma = 5.7 \times 10^{-4}$ mS/cm$^2$ is the tonic GABA action potential scaling factor, $f_b = 0.1$ Hz is the average basal network spiking frequency at rest, $\omega = 0.004$ Hz is the GABA pump rate, and $g_{basal} = 0.1$ mS/cm$^2$ is the value at which GABA transporters reverse.[14] The variable $f_m(t)$ is the average firing rate across all cells in the population grid and is described the equations:

$$f_m(t) = \frac{1}{M}\sum_{i=1}^{M} f_i(t)$$

$$\frac{df_i(t)}{dt} = \sigma\left(\frac{N(t)}{W} - f_i(t)\right)$$

Where $f_i(t)$ denotes the slow-activating firing rate function of neuron $i$ in the populational grid, W = 2 seconds is the time window, and N(t) is the number of spikes over the time range W, t, and $\sigma = 1$ (1/s) is the firing rate time factor.

And for the dendritic synaptic currents, $I_{syn,d}$:

$$I_{syn,d} = I_{GABA-B} + I_{AMPA}$$

Where

$$I_{GABA-B} = g_{GABA-B}(V_{m,d} - E_K)$$

$$I_{Glu,D} = g_{Glu}(V_{m,d} - E_{Glu})$$

The dynamics of GABA conductivity is described by

$$g_{GABA-A} = \bar{g}_{GABA-A}\left(1 - \exp\left(\frac{-(t-t')}{\tau_1}\right)\right)\exp\left(\frac{-(t-t')}{\tau_2}\right)$$

Where $\bar{g}_{GABA-A}$ (in mS/cm2) is the maximal conductivity of the GABA-A receptor, t (seconds) is the current time, t' (seconds) is the time of the most recent somatic action potential, and $\tau_1, \tau_2$ (seconds) is the decay constant. For this model, we set $\tau_1 = 0.002$ and $\tau_2 = 0.05$ .

The glutamatergic conductance dynamics were represented using time-dependent synaptic kernels, following earlier conductance-based models.[7,16,17]

$$g_{Glu} = g_{Glu,\text{int}} + g_{Glu,ext}$$

$$g_{Glu,\text{int}} = \frac{1}{M}\sum_{i=1}^{M}\bar{g}_{Glu}\left(1 - \exp\left(\frac{-(t-t_{spike,i})}{\tau_{s,1}}\right)\right)\exp\left(\frac{-(t-t_{spike,i})}{\tau_{s,2}}\right)$$

$$g_{Glu,ext} = \bar{g}_{Glu}\left(1 - \exp\left(\frac{-(t-t_{Glu,i})}{\tau_{s,1}}\right)\right)\exp\left(\frac{-(t-t_{Glu,i})}{\tau_{s,2}}\right)$$

Where M is the number of neurons synaptically connected to the target neuron, $\bar{g}_{Glu}$ (in mS/cm²) denotes the maximal glutamate-receptor conductance, $t_{spike,i}$ denotes the time (s) of the most recent glutamatergic spike from neuron i, $t_{Glu,i}$ denotes the time (s) of the most recent external glutamatergic spike following a Poisson process, and $\tau_{s,1}$ and $\tau_{s,2}$ denote the rise and decay constants of the glutamate receptor and were set to $\tau_{s,1} = 3 \times 10^{-4}$ s and $\tau_{s,2} = 3 \times 10^{-3}$ s.

### Supplementary Table 1. Conductance parameters

| Parameters | Value (mS/cm$^2$) | Reference |
|---|---|---|
| $\bar{g}_{leak,s}$ | 0.1 | 8 |
| $\bar{g}_{leak,d}$ | 0.025 | 12 |
| $\bar{g}_{Na}$ | 30 | 8 |
| $\bar{g}_{K}$ | 15 | 8 |
| $\bar{g}_{K-P}$ | 0.077 | 12 |
| $\bar{g}_{Na-P}$ | 0.022 | 12 |
| $\bar{g}_{Ca}$ | 0.15 | 18 |

### Supplementary Table 2. Equilibrium potentials

| **Parameter** | **Value (mv)** | Reference |
|---|---|---|
| $E_K$ | -85 | [8*] |
| $E_{Na}$ | 50 | [8*] |
| $E_{leak}$ | -70 | [8*] |
| $E_{Cl}$ | -70 | [16] |
| $E_{GABA}$ | -70 | [15] |
| $E_{Glu}$ | 0 | [16] |
| $E_{Ca}$ | 70 | [19] |

* Values adapted to be consistent with a resting potential of −70 mV

### Supplementary Methods 6. Network and extracellular-potential model

The network consisted of a 1,000 × 250 × 500 µm volume represented computationally as a 20 × 2 × 10 grid, as described previously.[7] The x–z plane represented individual neurons, whereas the y axis represented soma and dendrite compartments. Specifically, j = 1 corresponded to the soma and j = 2 to the dendrite for each neuron in the x–z plane.

For the soma, the injected current was set to −0.44 µA/cm$^2$ with fluctuations from a synaptic-noise term and a neuron-specific modulatory recovery current motivated by the spike-history-dependent elevation of action-potential threshold observed in vivo.[20] The recovery current depended on the timing of the most recent action potential and followed the dynamics given below:

$$\frac{dI_{curr}(x,t)}{dt} = \rho(\mathrm{N_s}(\mathrm{x,t})\mathrm{r_{curr}} - I_{curr}(x,t))$$

Where $I_{curr}(x,t)$ is the recovery current at time $t$ for neuron $x$ in the grid, $\mathrm{N_s}(\mathrm{x,t})$ is the number of spikes over the past second from time $t$ for neuron $x$, $\rho$ is the recovery current decay constant, and $\mathrm{r_{curr}}$ is a scaling constant for the strength of the recovery current dependent on the number of previous spikes. For this model, $\rho = 0.01$ and $\mathrm{r_{curr}} = 0.02$.

The injected current for dendrites was set to 0.8 µA/cm$^2$, consistent with the current range used to sustain subthreshold membrane-potential oscillations.[12]

The coupled equations are iterated over the grid using a forward Euler numerical method, with a time step of 0.1 (in units of milliseconds).

Extracellular potential at a recording site was calculated from a distance-weighted summation of transmembrane currents following established field-potential and ephaptic-coupling approaches.[21-24] The extracellular potential at grid point z was given by

$$V_{e,k}(z,\ t) = SF * \frac{\rho}{4\pi} \sum_{x\,\epsilon\,P} \left( \frac{I_s(x,t)}{r_s(x,z)} + \frac{I_d(x,t)}{r_d(x,z)} \right)$$

Here SF is the stacking factor; extracellular resistivity and extracellular potential are defined for the soma and dendrite compartments, and the set includes all neurons in the population grid. To avoid singularities, neurons occupying the same position as the recording point were omitted from the summation. The extracellular resistivity followed the value used in the immediate physiological implementation.[24] The somatic and dendritic transmembrane currents were then calculated as shown below.

$$I_s(x,t) = A_s\left(I_{ion,s} + I_{syn,s} + I_{res,s} - I_{inj,s}\right)$$

$$I_d(x,t) = A_d\left(I_{ion,d} + I_{syn,d} + I_{res,d} - I_{inj,d}\right)$$

Somatic area was based on a 20-µm-diameter soma, and dendritic area was taken from published morpho-electrotonic estimates.[25,26]

Distances were calculated as Euclidean distances between each neuronal compartment and the recording position.

For the two-compartment model, $dV_{e,s} := V_{e,s}(z,\ t) - V_{e,s}(z,\ t-dt)$ and $dV_{e,d} := V_{e,d}(z,\ t) - V_{e,d}(z,\ t-dt)$.

The stacking factor represents the number of biological neurons represented by each grid point.[21] Using an approximate cortical neuronal cellular density of 57,000 cells/mm3 and the grid-step dimensions, SF was set to 35.625.[27]

Model pharmacological manipulations. For the normal resting-state condition, we set $\bar{g}_{Glu} = 0.15$ mS/cm$^2$, $g_{GABA-A} = 0.3$ mS/cm$^2$, and $\alpha_{GABA} = 1$. To simulate DGG, we retained the remaining resting-state parameters and set the glutamate conductance to $\bar{g}_{Glu} = 0$ mS/cm$^2$. To simulate muscimol after DGG, we retained the DGG parameters and set $\alpha_{GABA} = 2$. For PTX, we retained the resting-state parameters but set $g_{GABA-A} = 0$ mS/cm$^2$ and the tonic GABA scaling parameter to $\alpha_{GABA} = 0.1$.

In simulations comparing models with and without extracellular-potential feedback, extracellular fields had minimal effect during the resting state with normal synaptic connectivity. After glutamate conductance was removed, however, extracellular-potential feedback increased nearby neuronal recruitment, sharpened synchrony and increased the amplitude and delineation of compound LFP burst peaks. These simulations were used to test a plausible local recruitment route; the availability-reservoir control law itself does not require extracellular coupling as the exclusive mechanism.

### Supplementary Note 1. Truncated waiting-time comparison between DGG and picrotoxin

To compare next-burst timing between DGG and picrotoxin on a common support, suppression durations were analysed over the range 0.5–2.5 s. The lower bound of 0.5 s was the operational cutoff for defining a suppression period. For visualization and fitting, the support was shifted so that 0.5 s corresponded to zero. The mean suppression duration over this range was 1.17 s for DGG and 1.37 s for picrotoxin, giving shifted-exponential parameters lambda = 1.50 and lambda = 1.16, respectively (using the reciprocal of the interval mean to determine the lambda coefficient). DGG showed smoother exponential agreement and better exponential quantile agreement than picrotoxin. Log–log analysis showed no comparable linear scaling for DGG, whereas picrotoxin showed a heavier-tailed relation consistent with power-law structure. Maximum-likelihood estimation (in accordance to the methodology from [28]) using an x_min = 0.1 seconds (after shifting the interval by 500 ms), over the 0.5-2.5s interval, yielded alpha = 1.5, close to values reported for epileptic inter-event timing.[29] A separate descriptive regressor was run to visually show the linear relationship in the log-log panel representing the power law of the PTX suppression duration distribution. These analyses support a stochastic post-recovery component of DGG next-burst initiation, whereas picrotoxin recurrence reflects structured disinhibited propagation.
